# Extinction Risk of Sonoran Desert Annuals Following Potential Changes in Precipitation Regimes

**DOI:** 10.1101/2022.02.02.478887

**Authors:** William S. Cuello, Sebastian J. Schreiber, Jennifer R. Gremer, D. Lawrence Venable, Pete C. Trimmer, Andrew Sih

## Abstract

Rapid environmental change can affect both the mean and variability in environmental conditions. Natural selection tends to favour those organisms that best respond to such changes. Here, we consider delayed germination as bet hedging strategies for 10 Sonoran Desert annuals. We use a germination model parameterized with long-term demographic and climate data to explore potential effects of changes in the mean and variance in precipitation on the evolution of germination strategies, as well as the risk of extinction. We then explored the potential for evolutionary rescue in response to these changes. As expected, results indicate that as rainfall declines, or uncertainty in rainfall increases, all species have higher extinction risk (the former being more detrimental). These shifts also increased the benefit of delayed germination. Results also indicate that evolutionary rescue can often occur for small shifts, especially for more variable rainfall regimes, but would not likely save populations experiencing larger environmental changes. Finally, we identified life history traits and functional responses to precipitation that were most strongly correlated to the ability to cope with changes in rainfall and with potential for evolutionary rescue: dormant seed survivorship and, to a smaller degree, chance of reproduction and seed yield sensitivity to precipitation.

## Introduction

Current, ongoing, and future climate change has increased the need to assess species’ extinction risks in the face of change (Ceballos and Ehrlich, 2002; Barnosky et al., 2011; Urban, 2015; Change, 2007). Depending on the severity of future climatic shifts, anywhere from 0% to 54% of species could risk extinction if forecasts for precipitation and temperature hold (Urban, 2015; Foden et al., 2013; Miles et al., 2004). Understanding which species are more vulnerable and why is critical for assessing and predicting responses to climate change (Beaumont and Hughes, 2002). Although some species have shown the propensity for rapid evolutionary and plastic responses (Franks et al., 2007; Gonzalo-Turpin and Hazard, 2009; Franks et al., 2014; M.L.Avolio and Smith, 2013; Avolio et al., 2013), others have shown an inability to adapt fast enough to climate change (Gonzalo-Turpin and Hazard, 2009; Franks et al., 2014; Bush et al., 2016; Matz et al., 2018). Understanding to what extent individual species are already affected by current climate regimes, as well as future responses, reveals ecological and evolutionary responses to shifting climates, as well as providing information for predicting and mitigating effects of change.

Of course, some species have always contended with unpredictably varying environments (Stabeno et al., 2001; Mantua et al., 1997; Bernhardt et al., 2020). In these environments, populations are expected to evolve bet-hedging strategies, i.e., strategies that minimize between-year variance in fitness at the expense of having lower fitness in some years (Menu and Debouzie, 1993; Danforth, 1999; Cohen, 1966; Ellner et al., 1999; Seger, 1987; Ellner, 1987). In doing so, environmental risk can be spread across individuals from the same lineage and across multiple years, so that they may survive catastrophic years (Danforth, 1999; Venable, 2007). For example, plant species in the Mediterranean, the desert, and other extreme and variable environments rely on delayed germination through seed dormancy and seed persistence in order to buffer against disastrous years (Venable, 2007; Koornneef et al., 2002; Klupczyńska and Pawlowski, 2021; Tielborger et al., 2012). Here, seed dormancy refers to the prevention of intact, viable seeds from germinating under favorable conditions (Venable, 2007; Allen and Meyer, 2002; Carta et al., 2013). We take seed persistence to be the survival of seeds in the soil seed bank for ≥ 1 year (Walck et al., 2005). In essence, these traits minimize risk of extinction by hedging their bets against uncertainty.

How well bet hedging traits will work under future climate regimes is the focus of increasing attention. Organisms’ ability to cope with and persist in these regimes depends on how well their previously adaptive strategies function in the new conditions (Avolio et al., 2013; Sih et al., 2011; Sih, 2013; Crowley et al., 2019; Tielbörger et al., 2014), as well as how rapidly species can evolve traits in response to change, and how effective adaptation is to this change (Franks et al., 2014; Franks and Weis, 2008; Sultan et al., 2013; Anderson and Song, 2020). For example, annuals and perennials have been shown to evolve flowering time in response to heat and drought stress or frost damage (Franks et al., 2007; Franks and Weis, 2008; Miller-Rushing and Primack, 2008; Rhoné et al., 2010; Colautti and Barrett, 2013). This evolutionary adaptation has even been observed on short time scales (Franks et al., 2014; Nevo et al., 2012; Anderson et al., 2012; Franks et al., 2016). In fact, the flowering time of *Boechera stricta* (Brassicaeae), a mustard native to the US Rocky Mountains, has advanced significantly by 0.34 days per year between 1973 and 2011 due to warmer temperatures and earlier snowmelt dates (Franks et al., 2014; Anderson et al., 2012). Desert annuals have also been shown to evolve seed dormancy fractions in response to climatic variation and may do so on short time scales (Klupczyńska and Pawłowski, 2021; Clauss and Venable, 2000; Volis and Bohrer, 2013; Fernández-Pascual et al., 2013). However, while some populations and species have adjusted well to environmental change, evidence of climate maladaptation are growing (Franks et al., 2014; Crowley et al., 2019; Anderson and Song, 2020; Potvin and Tousignant, 1996; Springate et al., 2011; Robertson et al., 2013; Wilczek et al., 2014; Anderson and Wadgymar, 2020).

To explore the potential impact of climate change on species that inhabit a highly variable desert community experiencing substantial climate change (Kimball et al., 2010; Huxman et al., 2013), we turn to the winter annuals of the Sonoran Desert. In this desert, water (or lack thereof) is a key driver for trait evolution and population dynamics. Here, annual plants must contend with low and variable rainfall and high and variable temperature (Cox et al., 1988; Búrquez et al., 1999; Gremer et al., 2012). Indeed, this community has become a model system for understanding bet hedging through delayed germination in a variable environment (Venable, 2007; Gremer and Venable, 2014; Gremer et al., 2016; Cuello et al., 2019). Although species in this community have evolved delayed germination to buffer populations against unfavorable years under historical precipitation regimes (Venable, 2007; Gremer and Venable, 2014), it is unclear whether they will be susceptible to extinction when faced with more variable or overall less rainfall. Understanding this has become more and more imperative as climate change models predict that the Southwestern region of the United States will experience an increase in variance and a decrease in overall mean precipitation in upcoming decades (Brusca et al., 2013; Pendergrass et al., 2017; Dominguez et al., 2012). Indeed, warming and drying over the past 25 years in the Sonoran Desert has already corresponded with a shift in the species composition of winter annuals due to the delayed onset of germination-triggering winter rains (Kimball et al., 2010). Thus, it is also important to understand how species-specific precipitation responses and life-history traits buffer or hinder population densities with shifts in precipitation regimes.

Here, we explore how Sonoran Desert annuals’ populations will be influenced by predicted climate changes, particularly shifts in the mean and variance of precipitation (Groisman and Knight, 2008; James and Washington, 2013; Trenberth et al., 2014). Long-term observations and simulations show a trend of decreasing precipitation in the US Southwest (Kimball et al., 2010; Huxman et al., 2013; MacDonald, 2010); specifically, precipitation decreased by 6.6mm yr^*−*1^ in the winter annual growing season at the Desert Laboratory in Tucson, Arizona (Kimball et al., 2010). Such changes may have strong effects on these annuals, particularly in terms of affecting water availability (Kimball et al., 2010; Huxman et al., 2013; Angert et al., 2010). Although this can be further exacerbated by high temperatures, increased evapotranspiration, decreased runoff, and increased drought (MacDonald, 2010; Huggins et al., 2010; Bradford et al., 2020; Dai, 2011), we focus on understanding how reductions in mean precipitation and increased variability in precipitation, in particular, affect population densities of desert annuals. We then ask how these effects might be mediated by their evolved life history and bet-hedging strategies. Will rapid evolution towards new optima in new regimes help populations recover from the negative effects of climate change on population viability? Which life history traits (e.g., seed survival rates) or functional responses to precipitation have more impact on the ability to cope with changes in rainfall and with potential for evolutionary rescue (i.e., the ability to recover in population density through adaptation)?

To answer these questions, we use a 30-year data set for 10 species of Sonoran Desert annual plants to parameterize population-dynamic models for each species and calculate their evolutionarily stable strategies for germination fractions (ESS germination fractions) (Cuello et al., 2019; Venable). We determine population size distributions given by the models under the historical precipitation regime. To examine the relative vulnerability of each species to precipitation shifts, we identify species-specific population density thresholds as a metric of extinction risk. To understand the ability of each species to cope (or not) with changes in precipitation regimes using their previously adaptive germination strategies, we calculate the percentage of time population densities are projected to spend below these thresholds using their previously adaptive germination strategies. To assess their scope for evolutionary rescue, we calculate the percentage of time populations spend below this threshold, if they are allowed to rapidly evolve to the ESS germination fraction. Finally, we evaluate the extent to which these results can be attributed to species-specific functional responses or life history traits.

## Methods

Long-term demographic monitoring of the Sonoran Desert winter annual community has been conducted since 1982 at the University of Arizona’s Desert Laboratory at Tumamoc Hill in Tucson, AZ (see Appendix from Cuello et al. (2019)). Briefly, each year 72 plots are visited after germination-triggering rain events and individuals are mapped upon germination and followed for subsequent survival and fecundity (Venable, 2007; Gremer and Venable, 2014; Pake and Venable, 1996). The density of viable, non-germinating seeds have also been monitored every year since the 1989-1990 growing season (Venable, 2007; Gremer and Venable, 2014; Pake and Venable, 1996). From these data we calculated germination fractions and per-capita survival and reproduction for each species in the desert annual plant community. Here we focus on 10 common, abundant species as in previous studies (Venable, 2007; Gremer and Venable, 2014; Cuello et al., 2019).

Using the first 30 years (1984-2013) of precipitation and yield data, we created a precipitation-driven, bet-hedging model that tracked the annual seed densities of these 10 species (Cuello et al., 2019). This model incorporates seed yield as a function of total precipitation for each year, seed dormancy, and intraspecific competition as density-dependent feedback. Using this model, we now incorporate scenarios expected under future precipitation regimes and explore its impacts on population dynamics and evolutionary stable strategies (ESS) for germination. We analyze the extent to which evolutionary change can buffer populations from extinction. To do so, we simulate population dynamics under new precipitation distributions. We then identify each species’ ESS for germination fractions, which is the germination strategy that resists invasion attempts by any mutant population playing a different strategy, and compare these new ESS germination fractions with those of the historical precipitation regime.

To explore relative vulnerability to extinction under current and future precipitation scenarios, we specified a density threshold under which populations are likely to go extinct for each species. The species specific thresholds were quantified using densities from simulations of the population models under historical rainfall regimes. To assess change in potential risk following a climatic shift, we calculated the frequency at which populations fell below this historical threshold, which we called the percent time below (PTB) the historical threshold. Here, we consider higher PTB as higher potential for quasi-extinction risk and consider a species to be quasi-extinct if it reaches 100% PTB.

We explored new precipitation regimes with either reduced means or increased variances. For each of these precipitation regimes, we calculated the PTB given the species’ ESS germination fractions under the historical rainfall regimes. To understand how rapid evolution to the new ESS germination fractions reduce extinction risk, we calculated the corresponding PTB for species with the ESS for the new precipitation regime. We define the potential for evolutionary rescue to be the difference between these two percentages (the PTB with adaptation subtracted from the PTB without adaptation). This difference was always positive indicating that rapid evolution always reduced extinction risk. Finally, we statistically tested how species-specific life-history traits and functional responses to precipitation affected ESS germination fractions and how these, in turn, affected potential for evolutionary rescue. These statistical relationships allow us to identify which species traits determine greater vulnerability to extinction risk due to unfavorable precipitation shifts and their potential to reduce this risk by evolving.

### The Model

Let *n*_*t*_ be the density of seeds in the seed bank in year *t* for a species. A fraction of *g* seeds germinate each year. Under low density conditions, germinating seeds contribute *Y*_*t*_ seeds to the seed bank in year *t*; here *Y*_*t*_ is referred to as the low-density yield in year *t*. These seeds survive to the next year with probability *s*_fresh_ (i.e. the survival rate of fresh seeds). Negative density-dependence reduces yield by a factor 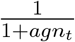, where *a* > 0 is a species-specific competition coefficient for the germinating population (Gremer and Venable, 2014; Cuello et al., 2019). A fraction of 1 − *g* seeds remain dormant until the next year, each surviving with probability *s*_dorm_ (i.e. the survival rate of dormant seeds) (see Figure 1A). Under these assumptions,

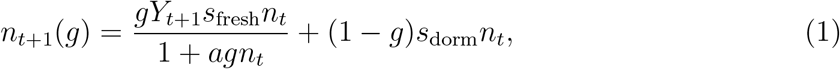

where the sequence *Y*_1_, *Y*_2_, … is independent and identically distributed. We used the measured survivorships, *s*_fresh_ and *s*_dorm_, and measured competition rates, *a*, from Gremer and Venable (Gremer and Venable, 2014). If the low-density *per capita* growth rate

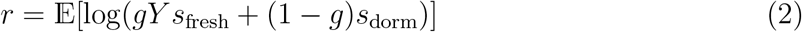

is negative, then *n*_*t*_ converges with probability one to zero. Otherwise, if *r* is positive, then *n*_*t*_ converges to a unique, positive stationary distribution 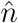 (Benaïm and Schreiber, 2009; Schreiber and Benaïm, 2019). We can approximate the stationary distribution of 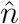 by one sufficiently long run of the model (Benaïm and Schreiber, 2009).

**Figure 1.**
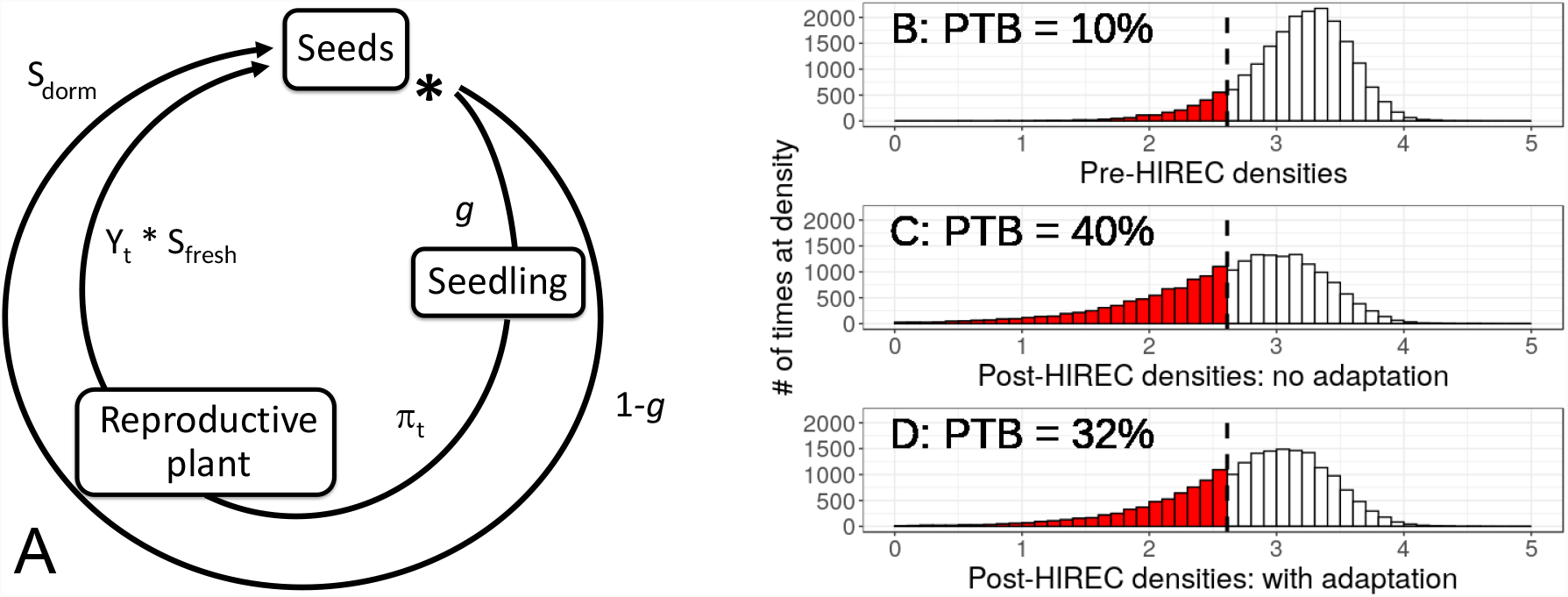
**Panel A**: the full life cycle diagram of a desert annual; an individual can either germinate with probability *g*, reproduce with probability *π*_*t*_, and produce fresh seeds *Y*_*t*_ of which a fraction *s*_fresh_ survives to the next year; otherwise, the seed stays dormant with probability 1 − *g* and survives with probability *s*_dorm_. **Panel B**: distribution of *Evax multicaulis* EVMU in the historical precipitation regime. **Panel C**: distribution of EVMU without adaptation after a 25% mean reduction in precipitation. **Panel D**: distribution of EVMU with adaptation after a 25% mean reduction in precipitation. In panels B through D, densities are on a log-scale; red indicates densities below the historical threshold, where the threshold is visually indicated by a vertical, black line.

To relate Eq.(1) to precipitation, we model low-density yield from precipitation, using 30 years of yield and precipitation data from 1984 to 2013 as done by Cuello et. al. (Cuello et al., 2019). Specifically, we fit a log-normal curve to the 30 years of measured precipitation data, gathered in Tucson, Arizona (Venable, 2007); let *P*_hist_ be a log-normal random variable with the same mean and variance as this fitted distribution, i.e.,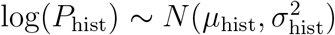, where *µ*_hist_ and 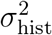 are the mean and variance of the log-transformed precipitation data. Following Cuello et. al. (Cuello et al., 2019), we model yield from precipitation via a two-step process, known as a hurdle model (Cuello et al., 2019; Mullahy, 1986). The first step is calculating the probability of reproductive success of a seedling *π*_*t*_:

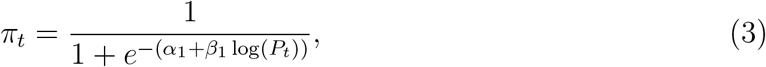

where the chance of reproduction on dry years *α*_1_ and sensitivity of chance of reproduction to precipitation *β*_1_ are species-specific parameters determined by a binomial regression relating the 30 years of low-density yield data to precipitation data (see Table 1). In particular, *α*_1_ determines the chance of reproduction for individuals when there is little precipitation, whereas *β*_1_ determines the extent to which this chance is increased for individuals when there are increased levels of precipitation.

**Table 1.**
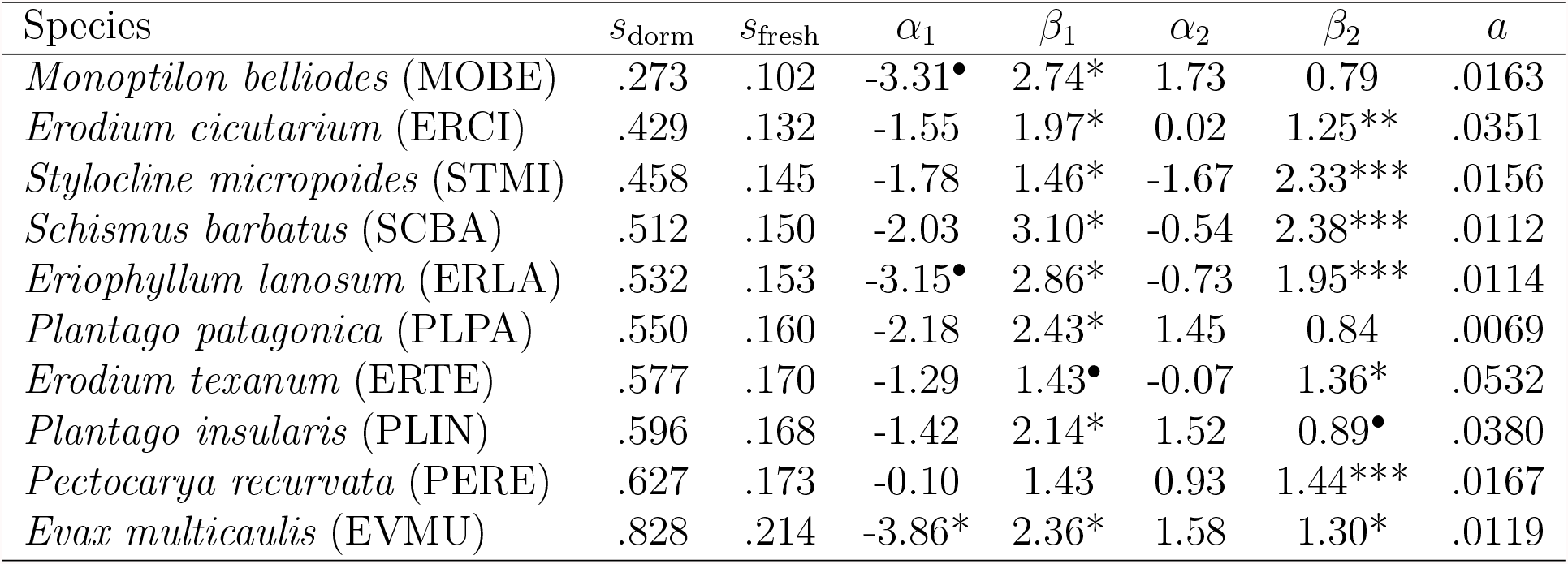
(Cuello et al., 2019): This table contains (from left to right) values for seed survivorships in the seed bank, seed survivorships of fresh seeds, chances of reproduction on dry years, sensitivities of chance of reproduction to precipitation, seed yield on dry years (upon successful reproduction), sensitivities of seed yield to precipitation, and intraspecific competition factors. Superscripted dots and stars next to *α* and *β* values indicate how significantly different these parameters are from 0: .1^•^, .05^*\**^, .01^*\*\**^, and .001^*\*\*\**^.

If an individual reproduces, i.e., clears the reproductive hurdle, its low-density yield *Y*_*t*+1_ equals (Cuello et al., 2019):

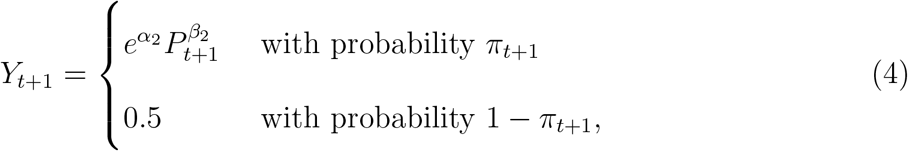

where *P*_*t*+1_ ∼ *P*_hist_. Here, seed yield on dry years (upon successful reproduction) *α*_2_ and sensitivity of seed yield to precipitation *β*_2_ are species-specific parameters determined by regressing measured, log-transformed, non-zero yield to precipitation (see Cuello et al. (2019); Table 1). In words, we first generate the probability of reproducing at all from year *t* to *t* +1, i.e., *π*_*t*+1_ (see Eq.(3)), which we can think of as clearing the “reproductive hurdle;” we then determine the number of fresh seeds that are produced *Y*_*t*+1_ (see Eq.(4)). Thus, variance in density values *n*_*t*_ are driven solely by variance in precipitation values *P*_*t*+1_.

For each species, we solved for the evolutionarily stable strategy for germination fraction (ESS value) under the historical precipitation regime *P*_hist_ (Cuello et al., 2019). To solve for an ESS value *g*, we consider a “mutant” population at very low density *ñ*, using a different germination strategy 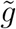. At low densities, this mutant population has negligible feedback on the resident population, but is influenced by the resident. Hence, we can initially approximate its dynamics by

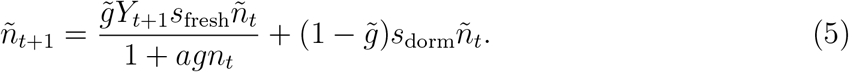

The success or failure of the invasion of the mutant sub-population is determined by its stochastic growth rate when rare:

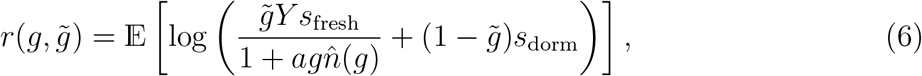

where 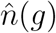 is the stationary distribution of the resident population. If 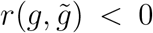, the invasion fails and if 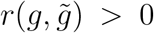, it succeeds (Schreiber and Benaïm, 2019; Chesson and Ellner, 1989). A germination strategy *g* is an ESS if 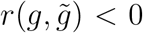 for all 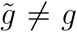, meaning no other strategy can invade when rare. One finds such an ESS germination fraction by solving for a 0 *< g <* 1 satisfying 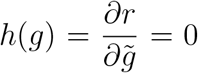; if this value exists, it is unique as 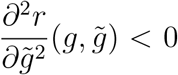 for all 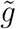 (Cuello et al., 2019). We denote the ESS germination fraction calculated under the historical precipitation regime, where each *P*_*t*+1_ is distributed as *P*_hist_ for all time *t* ≥ 0, as *ĝ*_hist_.

Since the initial (positive) density does not alter long-term population dynamics in this model, we choose *n*_0_ = 1 and iterate *n*_*t*_(*ĝ*_hist_) forward until it approximates the stationary distribution 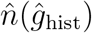 (Benaïm and Schreiber, 2009). Specifically, we take the last 90% of values from a run of 800, 000 time steps to estimate 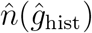. The 10 species’ long-term population density distributions are not similar, due to differences in their low-density yield distributions (Gremer and Venable, 2014).

To assess the amount of risk each species is facing in new precipitation regimes, we define a density threshold below which the population is considered at risk. To prevent bias against species with higher variance in their densities, we have species specific thresholds based on their population dynamics during the historical precipitation regime. Specifically, we define the historical threshold, *τ*_hist_, as the lowest decile of the stationary distribution of the population densities under the historical precipitation conditions 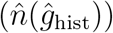, i.e., under one long run, the species spends approximately 10% of its time below *τ*_hist_ (see Figure 1B).

### Changing Precipitation Regimes

Using the historical threshold, we determine how a range of fixed changes to *P*_hist_ (i.e., reducing mean or increasing variance in precipitation), affects the percent time below the threshold. Since *P*_hist_ is log-normal with 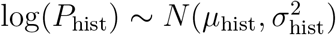,

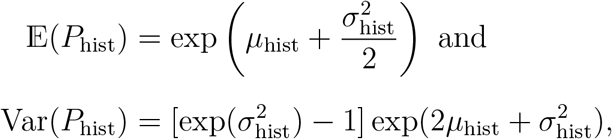

where 𝔼 (*X*) and Var(*X*) are the expected value and the variance of a random variable *X*, respectively.

To create a new precipitation distribution such that its standard deviation is equivalent to that of the historical regime (i.e., Var(*P*_new_) = Var(*P*_hist_)) and its mean is reduced by Δ_mean_*100%, we solve for a new mean *µ*_new_ and standard deviation *σ*_new_ such that 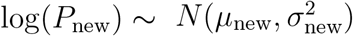, where

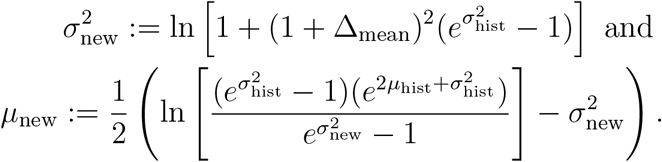

To create a new precipitation distribution whose mean is fixed (i.e., 𝔼 (*P*_new_)) = 𝔼 (*P*_hist_)), but standard deviation is increased by Δ_SD_ * 100%, we solve for a new mean, *µ*_new_, and standard deviation, *σ*_new_, such that 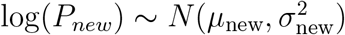. Namely,

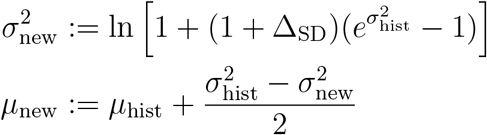

To determine an upper value to which we would increase standard deviation in precipitation, we used a relationship between rising temperatures on the global scale and increased variability (Pendergrass et al., 2017); we determined the range of standard deviation values by multiplying the percent change in standard deviation in precipitation per unit change in temperature 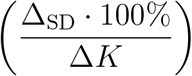, where Δ*K* is the change in Kelvin, to the forecasted increase in temperature per decade in the southwestern region of the United States (Pendergrass et al., 2017; Bai et al., 2014). Hence, we kept the percent increase in standard deviation between 0% and 25% (i.e. 0% ≤ Δ_SD_ · 100% ≤ 25%). For the sake of comparison, the interval of mean percent reduction in precipitation was also kept between 0% and 25% (i.e. 0% ≤ Δ_mean_ · 100% ≤ 25%); this upper value was consistent with predictions that mean precipitation would decrease in the southwestern United States over the next few decades by 20% (Dominguez et al., 2012)’s.

### Percent Time Below the Historical Threshold

For each new precipitation regime and each species, we calculated the percent time spent below the historical threshold (PTB) assuming no change in the species’ bet-hedging strategy (see Figure 1C). We note that for the non-native, naturalized species *Erodium cicutarium* (ERCI), its low-density stochastic growth rate was negative in the historical precipitation regime *P*_hist_. Consequently, its population densities asympotitically tended to 0 and, therefore, PTB has no meaning. Thus, we have omitted the species from our analyses and results.

To further contrast the relative resilience of the 9 species to climate change, we calculated the change in precipitation required to increase each species’ PTB by a standardized amount. A species is more resilient to climate change if (relative to another species) a larger decrease in mean rainfall or increase in year-year variation in rainfall is required to cause that species’ PTB to increase by the standardized amount. For this standardized amount, we choose the largest PTB (13%; see Figure 2) that all species exhibited over our changes in precipitation. As the species’ PTB responses to these changes (see Figure 2) generally did not cross, we get very similar rankings for relative species resilience regardless of our chosen values for the standardized increases in PTB.

**Figure 2.**
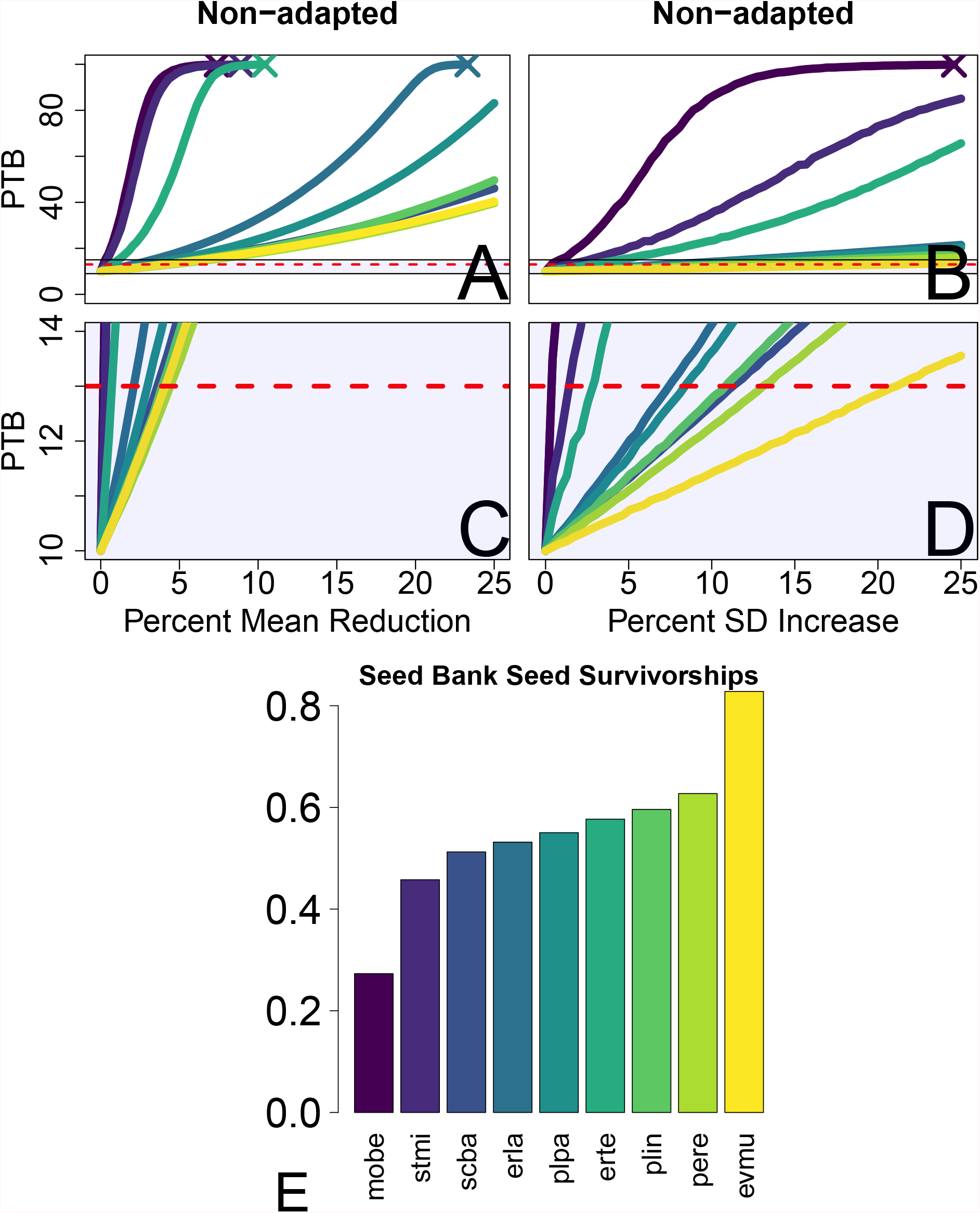
**First row, Panels A (left) and B (right)**: the percent time spent below the historical threshold (PTB) for each species as a function of percent reduction in mean precipitation (left) and increased variance in precipitation (right). The × markers designate 100% PTB or quasi-extinction. The dotted red line at 13% indicates the highest (rounded down) PTB realized by all species. **Second row, Panels C (left) and D (right)**: zoomed in images of translucent purple boxes featured in A (left) and B (right). **Third row, Panel E**: seed bank seed survivorships of the nine species from lowest (darkest) to highest (brightest).

### Adaptation to a New Precipitation Regime

For each new precipitation regime *P*_new_, we drew precipitation values *P*_*t*+1_ distributed as *P*_new_ and used these to generate corresponding *Y*_*t*+1_ values via Eq.(3). We then calculated the new ESS germination fraction *ĝ*_new_ in regime *P*_new_ using Eq.(6).

For each new precipitation regime and ESS germination fraction, we iterated *n*_*t*_(*ĝ*_new_) until the process had approximated the stationary distribution 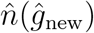 (see Figure 1D). Here, we calculated the PTB of 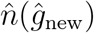, i.e., the percentage of time in which a species stayed below historical threshold *τ*_hist_ after adapting to the new ESS germination fraction (see Figure 1D; red bars). Finally, we calculated the *potential for evolutionary rescue* as the reduction in PTB due to adaptation to the new precipitation regime.

### Identifying the role of plant traits on extinction risk and potential for evolutionary rescue

We statistically analyzed the extent to which life history traits and functional responses drove species’ resilience and relative changes in their bet-hedging strategies in response to changes in precipitation. In particular, we calculated the minimum percentage reduction in mean rainfall or percentage increase in variance necessary to reach PTB = 13% (i.e., species’ resilience). For each species, we then calculated the relative reduction in ESS germination fractions from the historical regime to the first regime that made the species exhibit 13% PTB, i.e.,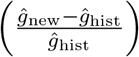. We then used AICc-selection and average models to predict species’ resilience (in both scenarios) and relative changes in ESS germination fractions (in both scenarios) from dormant seed survival rates (*ŝ*_dorm_), chance of reproduction on dry years 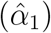, and sensitivities of seed yield to precipitation 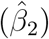 (Symonds and Moussalli, 2011; Burnham and Anderson, 2004; Burnham et al., 2011), as this life trait and these functional responses explain most of the variance in ESS germination fractions in the historical regime (Cuello et al., 2019). In total, 4 average models were created. Survival rates of fresh seeds *s*_fresh_, sensitivity of chance of reproduction to precipitation *β*_1_, and seed yield on dry years *α*_2_ were not used due to their statistical dependence with *s*_dorm_, *α*_1_ and *β*_2_, respectively (see Eq.(3) and Eq.(4)) (Cuello et al., 2019). Moreover, we did not include the strength of intraspecific competition *a* as a predictor, as nondimensionalization removes it from Eq.(1) (Cuello et al., 2019).

We then examined the statistical relationship between relative changes in ESS germination fractions and potential for evolutionary rescue in both scenarios. In particular, we calculated the potential for evolutionary rescue for species in the regime that first made species exhibit PTB = 13%. We then statistically tested for a monotonic relationship between relative changes in ESS germination fractions and potential for evolutionary rescue using Spearman Rank correlation tests. We used Fisher’s *Z* transformation to test for significance of the resulting Spearman Rank coefficients (Choi, 1977; Fieller et al., 1957).

## Results

As expected, percent time spent under the historical threshold (PTB) increased when we introduced climate change through reductions in mean precipitation and increased precipitation variation (Figure 2A,B). Decreases in mean precipitation of up to 25% had substantial negative effects on population persistence. Four species (MOBE, STMI, ERLA, and ERTE) went quasi-extinct (reached 100% PTB; see × marks in Figure 2) with 3 of those 4 going quasi-extinct with only a 5−10% reduction in mean precipitation. One species (PLPA) came relatively close to quasi-extinction (> 80% PTB), whereas the remaining 4 species (SCBA, PLIN, PERE, and EVMU) showed only moderate increases (from 30 − 50%) in PTB. In contrast, an increase of up to 25% in the variance in precipitation had smaller negative impacts. Only MOBE went quasi-extinct with a 25% increase in standard deviation. STMI came relatively close (> 80%) to quasi-extinction; ERTE experienced a moderate increase (> 60%); however, the remaining 6 species experienced very little change (increasing from 10% to 13 − 20%) in PTB. Whenever species went quasi-extinct or reached 100% PTB, their low-density stochastic growth rate was negative and their population densities declined exponentially. The most resilient species was EVMU (see Figure 2B,D; yellow line), thus making the largest (rounded down) PTB exhibited by all species across both scenarios 13% (see Figure 2; dotted red lines).

The leading driver for species’ resilience to both reduced rainfall and increased variance was dormant seed survivorship *s*_dorm_ (see rows 1 and 2 of Table 2). Species with higher dormant seed survivorship were much more resilient than other species in both scenarios. In reduced mean scenarios, higher chance of reproduction on dry years 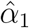 also boosted resilience but to a lesser degree. On the other hand, sensitivity of seed yield to precipitation 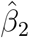 had a small but negative impact on resilience (see row 1 of Table 2). In the case of increased variation in rainfall, species with lower chance of reproduction on dry years were instead marginally more resilient to increased variation in rainfall; although, sensitivity of seed yield to precipitation still minimally decreased resilience (see row 2 of Table 2).

**Table 2:** The first and second lines are the average models for predicting the resilience of species against reduced rainfall or increased variance in rainfall, i.e., the minimum percentage decrease in mean and minimum percentage increase in SD, respectively, to realize PTB = 13%. The third and fourth lines are the average models for predicting relative changes in ESS germination fractions from the historical regime to the one that first produced PTB = 13% in the reduced mean scenario and the increased standard deviation (SD) scenario, respectively. Here, the standardized predictors are seed bank seed survivorship (*ŝ*_dorm_), chance of reproduction on dry years 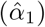, and sensitivity of seed yield to precipitation 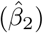. The average model coefficient for each predictor is calculated using the weighted sum of that predictor’s coefficient in each AICc model (see 3 to 6 found in Appendix A); here, the weight is the Akaike weight of the corresponding model.

| Response variable | Intercept | $\hat{s}_{\text{dorm}}$ | $\hat{\alpha}_1$ | $\hat{\beta}_2$ |
| --- | --- | --- | --- | --- |
| (Resilience)<sub>mean</sub> | 2.80 | 1.45 | 0.07 | -0.11 |
| (Resilience)<sub>SD</sub> | 8.18 | 5.21 | -0.29 | -0.03 |
| (Relative Reduction in ESS)<sub>mean</sub> | 1.38 | 0.58 | 0.01 | -0.03 |
| (Relative Reduction in ESS)<sub>SD</sub> | 2.58 | 1.32 | -0.06 | 0.13 |

ESS germination fractions decreased by minimal amounts in both reduced mean and increased variance in precipitation scenarios. The same four species that went quasi-extinct with reductions in mean precipitation experienced a very brief but steeper descent in ESS germination fractions when the reductions in mean precipitation approached their point of quasi-extinction (see Figure 3A). We attribute this to densities being so small that ESS germination fractions were equivalent to that of a population experiencing no intraspecific competition, i.e., finding the ESS of Eq.(1) with 0 *< n*_*t*_ *<<* 1. Results were similar for species in the increased variance in precipitation scenario, although they decreased by an even smaller than that of the reduced mean scenario (Figure 3B).

**Figure 3.**
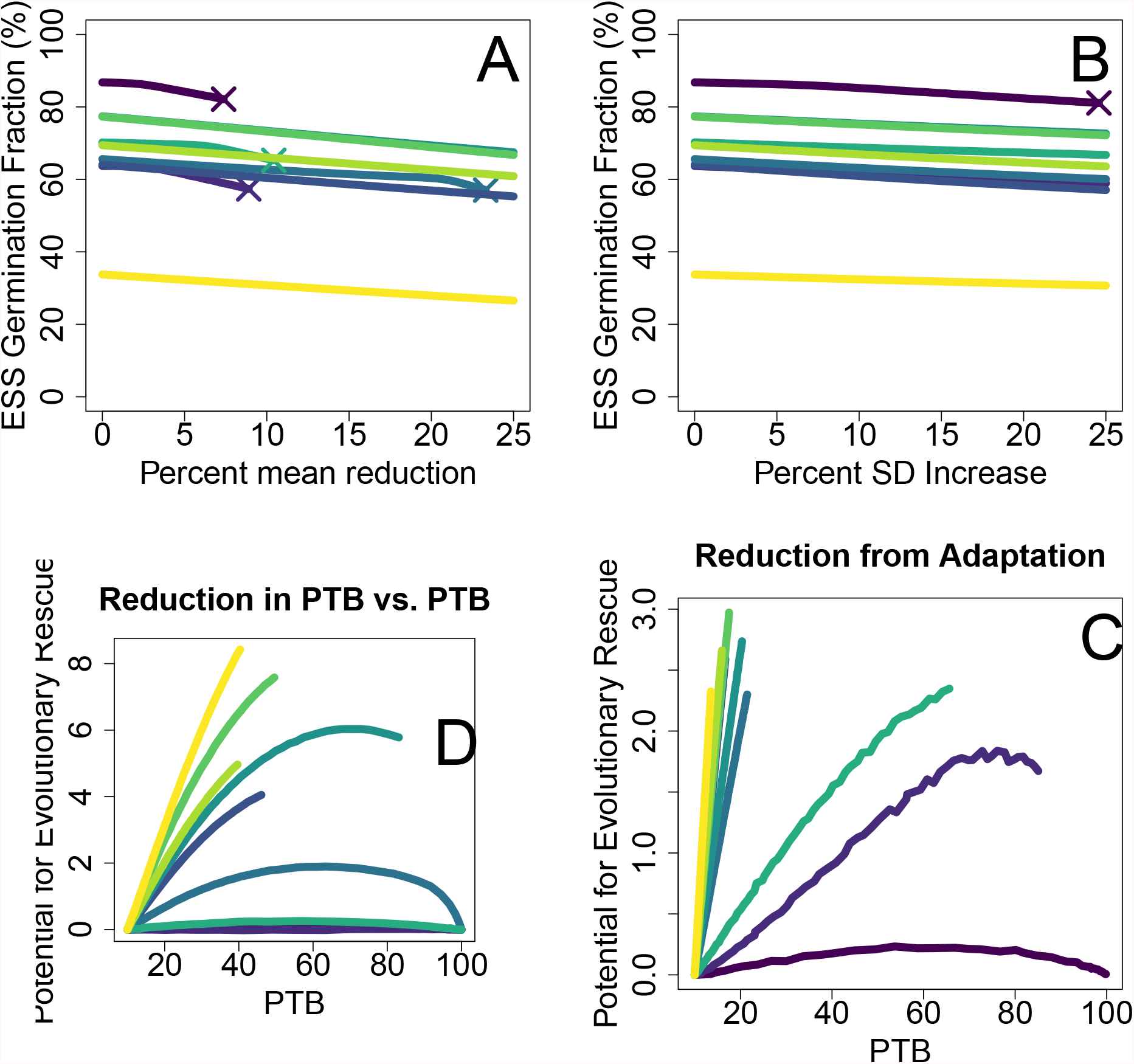
**First row, panels A (left) and B (right)**: panels A and B show how evolutionarily stable strategies for germination fractions (converted to percentages) change with reductions in mean and increases in standard deviation, respectively. The × mark at the end of ESS curves mark stochastic extinction. **Second row, panels C (left) and D (right)**: panels C and D show how potential for evolutionary rescue changes with respect to increases in percentage time below the historical threshold, due to reductions in mean precipitation and changes in standard in deviation, respectively.

Dormant seed survivorship was the most influential in driving the evolution of bet-hedging after climate change. In particular, species with higher dormant seed survivorship exhibited larger shifts in their ESS than other species (see Table 2). This prediction held for both responses to reduced precipitation and to increased variation in precipitation. In contrast, the other two life history traits, chance of reproduction in dry years 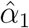 and sensitivity of seed yield to precipitation 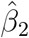, had little effect on the ESS germination fractions in rainfall regimes of increased variability and minimal effect in regimes with overall less rainfall. Specifically, species that had a higher chance of reproduction on dry years and a lower sensitivity of seed yield to precipitation hedged their bets marginally more (germinated marginally less) in drier regimes (see row 3 of Table 2); in contrast, species with a lower chance of reproduction on dry years and a higher sensitivity of seed yield to precipitation hedged their bets slightly more in variable regimes (see row 4 of Table 2).

Comparing PTB for adapted and non-adapted populations, we found that all species had a positive, albeit, relatively small, potential for evolutionary rescue. In the case of reductions in mean precipitation, the range in the potential for evolutionary rescue via shifts in ESS germination fractions was from 0% to 9% reduction in PTB (see Figure 3C). For the four species that experienced marginal increases in PTB with reduced precipitation, the potential for evolutionary rescue strictly increased as a function of PTB. In contrast, the three species that went quasi-extinct with only relatively small reductions in precipitation showed essentially no potential for evolutionary rescue. Perhaps most interestingly, the two species that were moderately vulnerable to quasi-extinction exhibited a parabolic relationship in potential for evolutionary rescue. With low-moderate reductions in mean rainfall (causing a moderate-large increase in PTB), shifts in germination frequency increased their potential for some evolutionary rescue. Specifically, for the case of reductions in mean precipitation, we found a moderately strong, positive relationship (Spearman rank of 0.68; Fisher-transformed *z*-score of 1.973; *p* < .05) between relative change in ESS germination fractions and potential for evolutionary rescue (see Figure 4A). But when the increase in PTB exceeded about 70%, the potential for evolutionary rescue decreased.

**Figure 4.**
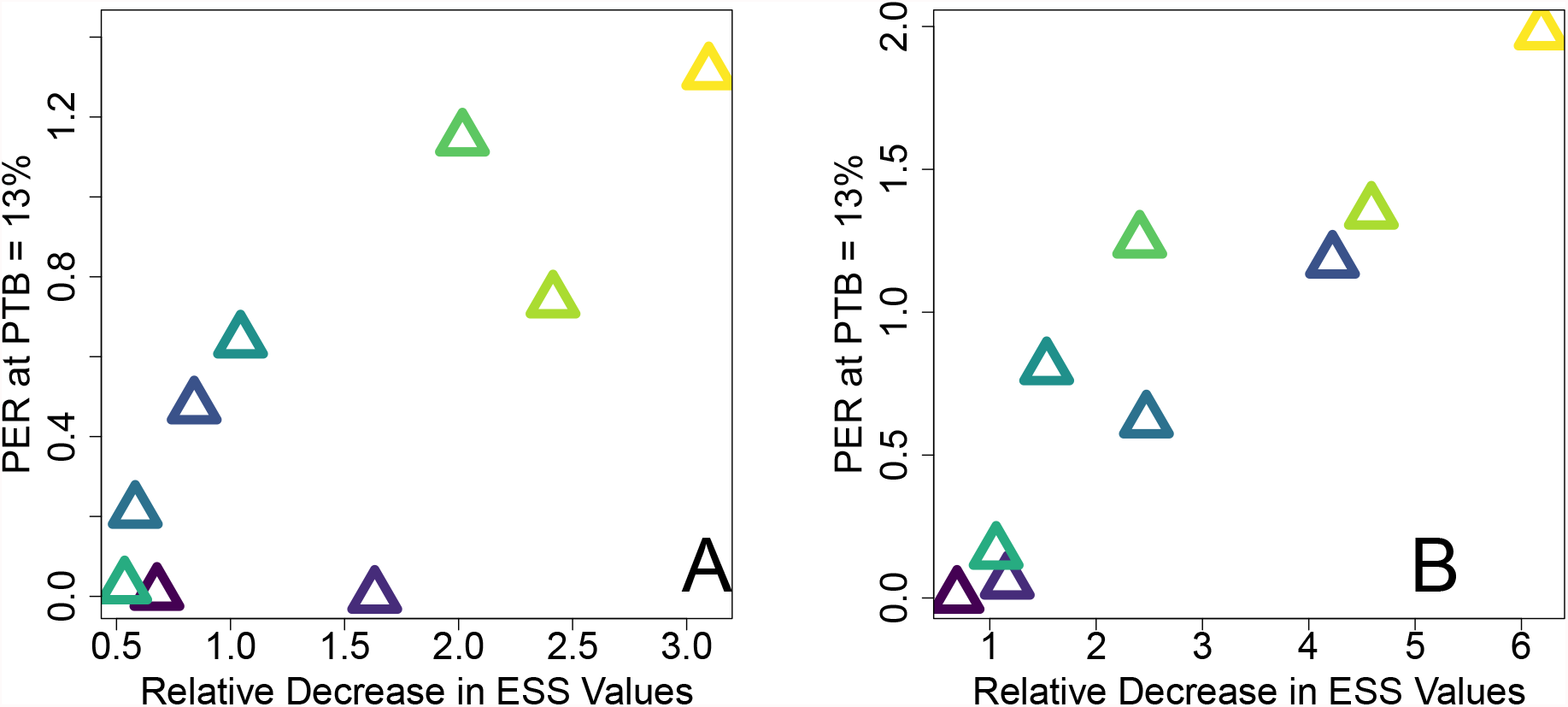
**Panel A (left)**: Potential for evolutionary rescue at PTB = 13% versus relative decrease in ESS from the historical ESS value, due to reduced mean in precipitation (Spearman correlation of 0.68). **Panel B (right)**: potential for evolutionary rescue at PTB = 13% versus relative decreases in ESS, due to increased variance in precipitation (Spearman correlation of 0.9).

In contrast, adaptation to the ESS germination fractions under increased variance resulted in a smaller range in potentials for evolutionary rescue: 0% to 3% reduction in PTB (see Figure 3D). *Monoptilon belliodes*, which went quasi-extinct, showed minimal potential for evolutionary rescue and particularly exhibited 0% potential for evolutionary rescue as it approached quasi-extinction. However, this is not to be interpreted as species adapting better to reduced mean scenarios. There was a lower risk of extinction for increased variance scenarios and, consequently, less PTB to recover through adaptation; there was also more potential evolution of germination strategies, and that evolution was more effective at reducing PTB under increased variance scenarios. We confirmed these latter statements through the Spearman rank test showing the stronger positive relationship (Spearman rank 0.9; Fisher-transformed *z*-score of 3.5; *p* < .001) between the relative change in the ESS and potential for evolutionary rescue (see Figure 4B) and ANOVA tests, showing that the initial slopes of the curves in Figure 3C were significantly less than those in Figure 3D (*p* < .001 for all nine species). As with the reduced mean scenario, species that came close to quasi-extinction also exhibited a parabolic relationship, its peak potential for evolutionary rescue being *<* 2% (after PTB exceeded 70%). The remaining seven species that experienced little to moderate increases in PTB from increased variance all strictly increased in potential for evolutionary rescue. We note, however, that species exhibiting a strict increase in potential for evolutionary rescue would not do so if we allowed for even worse regimes: we found that all species would eventually go quasi-extinct and eventually exhibit zero potential for evolutionary rescue with sufficiently large (> 25%) reductions in mean or increases in variation.

## Discussion

While shifts in variability are often expected to have stronger effects than changes in mean conditions (Easterling et al., 2000; McLaughlin et al., 2002; Vincenzi, 2014), for this community of annual plants in the Sonoran Desert, we found that decreases in mean precipitation were typically more detrimental to species than parallel increases in variance. In fact, three more species went extinct with overall less rainfall than increased uncertainty. At first this seems surprising, but these species already inhabit and are thus adapted to a highly variable environment (Venable et al., 1993; Shreve and Wiggins, 1964). In particular, bet-hedging strategies (e.g., seed dormancy) are typically employed by winter annuals to mitigate the risk of increased variance, rather than buffer against overall, decreased rainfall (Cohen, 1966; Venable, 2007; Philippi, 1993; Ellner, 1985; Cohen, 1968). Moreover, species supplement these bet-hedging strategies with species-specific functional responses to rainfall, some of them exhibiting a tradeoff of producing fewer seeds in drier years but capitalizing on high seed production in wet years (Angert et al., 2007; Huxman et al., 2008). However, such bet-hedging and response strategies become less and less relevant if little-to-no rain falls indefinitely, as seeds would then either eventually die or germinate to produce little-to-no yield.

Our results also indicate that species with higher dormant seed survivorships tend to stay relatively safe from detrimental precipitation shifts, especially in the case of increased variance. After all, populations can only gain the advantages of dormancy and germination if their seeds survive in the first place (Ooi, 2012). This advantage is, however, diminished in drier conditions: germination will typically lead to less seed yield and indefinite dormancy will eventually lead to the death of the seed. Indeed, *Evax multicaulis* and *Pectocarya recurvata*, the two species with the highest dormant seed survivorship, were the least affected by reduced mean precipitation and increased variance in precipitation, whereas *Monoptilon belliodes* and *Stylocline micropoides*, the two species with the two lowest dormant seed survivorships, except for the omitted *Erodium cicutarium*, experienced the highest reduction in population densities. In fact, seed survivorship may play a key role in differentiating the population dynamics of *Evax multicaulis* and *Stylocline micropoides* in more unfavorable regimes, as they historically have relatively higher demographic variances than the other observed species (Huxman et al., 2008).

To a lesser degree, chance of reproduction on dry years and sensitivity of seed yield to precipitation also determined how resilient species were to detrimental changes in both scenarios. In the reduced mean scenario, higher chance of reproduction on dry years and lower sensitivity of seed yield to precipitation buffered populations from overall less rain. To make sense of the latter, a lower sensitivity of seed yield to precipitation (or lesser ability to capitalize on wet-year bonanzas) is often the cost for species to be relatively more successful on dry years (Huxman et al., 2008; III et al., 1993; Angert et al., 2014). Thus, species better at capitalizing on large precipitation events often experience higher detriment when overall rainfall is decreased (Kimball et al., 2012). In the case of increased variability, our models indicate that it is more advantageous for annuals to suffer the cost of a lower chance of reproduction on dry years for a higher chance of reproduction on wet years. In other words, species must take advantage of the large pulses in rainfall that rarely accompany droughts rather than maintain a static chance of reproduction (Noy-Meir, 1973; Ault et al., 2016). In fact, it has been shown numerically that being able to capitalize on good conditions may sometimes be more beneficial than always performing moderately, depending on the pattern of variation in the environmental conditions and the fitness of the species as a function of these conditions (Liu et al., 2019).

This trade-off in resource uptake has already been observed for decades in the Sonoran Desert, as annuals have fallen on a spectrum of either being more stress tolerant in drought-like scenarios or having a higher ability to capitalize wet-year bonanzas. The former, which roughly produce little-to-moderate amounts of seeds regardless of precipitation regime, and are biologically characterized by low specific leaf area, high leaf nitrogen, and increased efficiency of photosynthetic processes related to light harvesting and performance in relatively cool conditions, are high water-use efficient (WUE) species (Gremer et al., 2012; Angert et al., 2007; Huxman et al., 2008). The latter, which roughly produce a large amount of seeds on wet years but little-to-no seeds on dry years, and are characterized by high specific leaf area, low leaf nitrogen concentration, and the ability to quickly respond to large precipitation pulses, are high rapid growth rate (RGR) species (Angert et al., 2007; Huxman et al., 2008). High-RGR species are favored in years with large, frequent rain events, whereas species with traits associated with high WUE do relatively better in years with infrequent rain and warmer temperatures at the end of the season (Kimball et al., 2012). Considering our results, we predict that unadapted annuals, already employing high-WUE strategies under the historical precipitation regime, will have a slight advantage in drier regimes. In contrast, we predict that unadapted annuals, already employing high-RGR strategies (particularly, ones that increase the chance of successfully reproducing on wet years) under the historical precipitation regime, will have slight, increased resilience against more variable regimes.

For the most part, species’ ESS germination fractions decreased very little in both the reduced mean rainfall and the increased variance in rainfall scenarios; although, species that decreased relatively more from their historical optimal values recovered the most through adaptation. This was, in part, determined by seed survivorship and functional responses to precipitation. If able to adapt to these new optima in drier conditions, annuals with high dormant seed survival rates would tend to germinate relatively less and consequently recover the most (though very little) through adaptation; here, it matters much less whether a species employs RGR or WUE strategies as these strategies had minimal effects on bet-hedging. For example, EVMU, with high dormant seed survivorship, can recover a slight amount through rapid adaptation in drought-like scenarios, whereas species like MOBE that sport a low dormant seed survivorship will most likely go extinct if no other mitigating strategies, such as seed dispersal, are employed (Venable and Brown, 1988). In contrast, we observed that desert annuals with higher seed survivorship and employing high-RGR strategies tend to hedge their bets more and recover the most (sometimes almost fully recovering from their losses) in more variable scenarios (Angert et al., 2007; Huxman et al., 2008; Gremer et al., 2013). We expect these species to stay relatively safe from low-to-mid range increases in variability with adaptation. Thus, rapid evolution is more effective for species such as *Evax multicaulis* that are able to capitalize on wet-year bonanzas (e.g., high-RGR species) and can keep their seeds alive and dormant for a relatively longer amount of time (Huxman et al., 2008). Although the potential for evolutionary rescue will vanish completely for all species if conditions become extremely variable or dry.

Of course, we are assuming rapid evolution, making it important to consider whether these species can adapt fast enough with respect to more deleterious regimes. This is especially important for species in more variable precipitation regimes, as high-seed-survivorship species (especially high-RGR species) are not only more resilient but are capable of recouping almost all their losses through adaptation, especially for small-to-moderate increases in variance. However, some plant species have only been partially successful in tracking recent warming (Corlett and Westcott, 2013; Jump and Penuelas, 2005). For example, successful recovery in density of plant communities in France significantly differed between lowland and highland forests in response to warming (Bertrand et al., 2011). On the other hand, desert annuals (e.g., the Californian *Brassica rapa*) have been shown to rapidly evolve other traits (e.g., flowering time) to cope with drought-like conditions (Sultan et al., 2013; Hamann et al., 2018; Dickman et al., 2019; Metz et al., 2020); such adaptations may give species enough time to adapt other traits, such as germination rates, to evolutionarily stable strategies, especially since evolution can occur on much shorter time-scales (e.g., decadal) than previously expected (Hamann et al., 2018; Hendry, 2016; Shaw and Etterson, 2012). Thus, a mix of phenotypic and evolutionary adaptation may help buffer such species until they successfully adapt their germination strategies to new regimes. To explore such effects of germination plasticity, one can incorporate environmental tracking into our seed density model and allow for year-to-year variance in germination fractions and assume positive correlation between germination fractions and precipitation. Moreover, we did not allow functional responses to precipitation to adapt to other precipitation regimes; modelers can also solve for ESS strategies for chance of reproduction on dry years and seed yield sensitivity to precipitation in altered regimes to see if the combined effects of adaptation in precipitation responses and germination rates further mitigate extinction risk.

Naturally, germination and bet-hedging go hand-in-hand with seed survivability, since the hedge is only as good as the safety it brings (Ooi, 2012). Particularly, we emphasize that seed survivability is the most important factor for both resilience to deleterious regimes and reaping the benefits of adaptation and may be the key to species’ survival in worse environments. For example, some grassland annuals have maintained their seed densities in spite of drought and delayed monsoon treatments, even if above-ground vegetation has been negatively impacted (Loydi and Collins, 2021). Others have seen their seed banks decline sharply in response to altered precipitation regimes (LaForgia et al., 2018). Thus, desert species exhibiting higher seed survivorship would likely have more time to evolve to optimal germination rates and might displace those that cannot keep up with climate change. In fact, it has been theoretically shown that certain species can exhibit slower rates of evolutionary response due to the presence of a seed bank, namely those whose germinating seeds spend (on average) a longer amount of time in the seed bank (Templeton and Levin, 1979). Such a species would be particularly susceptible to extinction if it exhibits a low seed survivorship as well. There are also more risks in the desert than uncertain or less rainfall, one example being seed predation. Seeds of some desert annuals may be at more risk than others due to increased granivore populations that prefer some seeds over others (Guo et al., 1998). Although there is clear evidence for positive frequency dependence in foraging patterns of granivorous rodents on winter annual seeds as a whole in the Sonoran Desert, certain species of plants, such as *Erodium texanum* and *Erodium cicutarium* were shown to have a higher probability of being harvested by granivores (Horst and Venable, 2017; Soholt, 1973). Yet the top predators that keep these granivores in check are already declining in population size due to increasingly less precipitation in this region (Brown et al., 1988; Iknayan and Beissinger, 2018; Riddell et al., 2019). Depending on what granivores are affected (i.e., generalists or specialists), this decline in top predators could uniformly increase consumption of seeds across all species or amplify predation on a subset of species. This would affect both species’ population levels and their potentials for evolutionary rescue and further alter the already-changing composition of Sonoran Desert annuals (Kimball et al., 2010). Hence, factors that impact seed persistence (and their relationships to one another) should be incorporated in future population models to numerically observe their effects on species’ densities.

As the effects of climate change are not restricted to shifts in precipitation regimes and predation, we also note that our model can be modified to incorporate other factors such as temperature. In an indirect manner, we capture some of the effects of temperature increase through our model; in particular, elevated evapotranspiration rates at higher temperatures is a large factor in there being overall less water within an environment and corresponds to reduced mean precipitation (Archer and Predick, 2008). However, we do not, for example, account for its effects on other traits such as seed survivorship. Excessively high soil temperatures may result in seed death (Ooi, 2012; Ooi et al., 2009). For a semi-arid species, *Wahlenbergia tumidifructa*, it was shown that seed longevity was reduced for those whose parent plants had grown under warmer temperatures (Kochanek et al., 2010). Thus, incorporating temperature into the model, as well as understanding its relationship to seed survivorship, life history traits, and seed yield could further our understanding of how plant species will cope with future precipitation regimes.

Our paper provides an example of how predictive frameworks can also be developed to understand the relation between species traits and their ability to cope with climate change (Newson et al., 2009; Pliscoff et al., 2014; Jones et al., 2013; Pereira et al., 2010), and specifically introduces a mathematical model for exploring the climatic impacts on bet-hedging annuals. Measuring dormant seed survivorship alone serves as a strong indicator on whether a species is buffered against uncertainty in, or overall less, rainfall. If similar precipitation, yield, and life history data can be aggregated for other desert annuals, our model could provide insight into whether those species are at risk to precipitation changes and have potential for evolutionary rescue. Moreover, accurately predicting future precipitation and climatic shifts in the upcoming decades will also provide valuable insight as to what species are more susceptible to endangerment or extinction in various environmental scenarios and whether climatic changes will be too rapid for adaptation in the first place (Pliscoff et al., 2014; Hoffmann and Sgrò, 2011). Such studies will be required for ecology to develop as a relevant predictive science in the 21st century.

## Appendix A: AICc-selection and Average Models

We calculated the AICc score of each model. We also found the *i*th model’s AIC difference Δ_*i*_ from the best model, its Akaike weight *w*_*i*_, its accumulated Akaike weight (acc *w*_*i*_), its evidence ratio (ER), the coefficients of each predictor (including the intercept term), and the adjusted-*R*^2^. The Akaike weights roughly correspond to the probability that a given model is the best approximating model; the evidence ratios provide measures of how much more likely the best model is than model *i* (Symonds and Moussalli, 2011).

We reported every model in increasing order of AICc scores (best to worst), rather than solely reporting the best model ; typically, models whose Δ_*i*_ *<* 2 should be kept in consideration for model selection (Symonds and Moussalli, 2011; Burnham and Anderson, 2004; Burnham et al., 2011). Here, Δ_*i*_ *<* 2 corresponds to models that are at least exp(−2*/*2) ≈ .36 times as probable to minimize information loss (Burnham and Anderson, 2002). In cases where there is at least one model whose Δ_*i*_ *<* 2, there are multiple pathways forward; since none of our best models had greater than 90% chance of the best approximating model, and there were typically two alternative models that satisfied Δ_*i*_ *<* 2, model uncertainty was evident (see Appendix; Tables 3 - 6) (Symonds and Moussalli, 2011). Thus, we took an approach based on all the models, and created an average model; in particular, to calculate the coefficient of each predictor in the average model, we took a weighted sum between the coefficients of each model multiplied to the corresponding Akaike weight *w*_*i*_ (Symonds and Moussalli, 2011). The purpose of this averaging is so that parameters with low Akaike weights will have little influence on the per-unit change in the response but will not be completely discounted (Symonds and Moussalli, 2011; Lukacs et al., 2010). We used this averaging, and the relative magnitudes of the predictors’ coefficients within each average model, to determine the extent to which the three predictors drove each response variable.

**Table 3:** all models for predicting the relative decrease in ESS from the historical precipitation to the ESS at 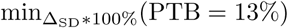; each uses a linear combination of at least one of the following predictors: 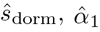, and 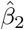. From left to right, we list AICc scores, the AIC difference from the best model (Δ*i*), the Akaike weight of the model (*w*_*i*_), the accumulated Akaike weight (acc *w*_*i*_), the evidence ratio (ER), the intercept term (L), and the coefficients of 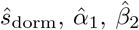, and the Adjusted-*R*^2^ of the model.

| Model | AICc | $\Delta_i$ | $w_i$ | acc $w_i$ | ER | $L$ | $\hat{s}_{\text{dorm}}$ | $\hat{\alpha}_1$ | $\hat{\beta}_2$ | Adj-$R^2$ |
| --- | --- | --- | --- | --- | --- | --- | --- | --- | --- | --- |
| $\hat{s}_{\text{dorm}}$ | 54.03 | 0.00 | 0.43 | 0.43 | 1.00 | 2.58 | 1.34 | 0.00 | 0.00 | 0.53 |
| $\hat{s}_{\text{dorm}} + \hat{\beta}_2$ | 55.10 | 1.06 | 0.25 | 0.68 | 1.70 | 2.57 | 1.35 | 0.00 | 0.35 | 0.50 |
| $\hat{s}_{\text{dorm}} + \hat{\alpha}_1$ | 55.78 | 1.74 | 0.18 | 0.87 | 2.39 | 2.58 | 1.35 | -0.19 | 0.00 | 0.46 |
| $\hat{s}_{\text{dorm}} + \hat{\alpha}_1 + \hat{\beta}_2$ | 56.72 | 2.68 | 0.11 | 0.98 | 3.82 | 2.56 | 1.36 | -0.21 | 0.36 | 0.43 |
| $\hat{\beta}_2$ | 61.62 | 7.60 | 0.01 | 0.99 | 44.50 | 2.69 | 0.00 | 0.00 | 0.32 | -0.10 |
| $\hat{\alpha}_1$ | 61.88 | 7.85 | 0.01 | 0.996 | 50.64 | 2.70 | 0.00 | -0.15 | 0.00 | -0.13 |
| $\hat{\alpha}_1 + \hat{\beta}_2$ | 63.53 | 9.50 | 0.004 | 1.00 | 115.32 | 2.68 | 0.00 | -0.17 | 0.33 | -0.27 |

### AICc Tables

**Table 4:** all models for predicting the 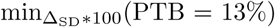; each uses a linear combination of at least one of the following predictors: 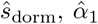, and 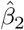. See Table 3 for variable interpretations.

| Model | AICc | $\Delta_i$ | $w_i$ | acc $w_i$ | ER | $L$ | $\hat{s}_{\text{dorm}}$ | $\hat{\alpha}_1$ | $\hat{\beta}_2$ | Adj-$R^2$ |
| --- | --- | --- | --- | --- | --- | --- | --- | --- | --- | --- |
| $\hat{s}_{\text{dorm}}$ | 71.93 | 0.00 | 0.47 | 0.47 | 1.00 | 8.20 | 5.22 | 0.00 | 0.00 | 0.71 |
| $\hat{s}_{\text{dorm}} + \hat{\alpha}_1$ | 73.19 | 1.26 | 0.25 | 0.73 | 1.88 | 8.15 | 5.25 | -0.84 | 0.00 | 0.69 |
| $\hat{s}_{\text{dorm}} + \hat{\beta}_2$ | 73.91 | 1.98 | 0.18 | 0.90 | 2.69 | 8.20 | 5.22 | 0.00 | -0.13 | 0.66 |
| $\hat{s}_{\text{dorm}} + \hat{\alpha}_1 + \hat{\beta}_2$ | 75.18 | 3.25 | 0.09 | 0.998 | 5.09 | 8.15 | 5.25 | -0.83 | -0.07 | 0.62 |
| $\hat{\alpha}_1$ | 84.11 | 12.18 | 0.001 | 0.999 | 440.60 | 8.63 | 0.00 | -0.69 | 0.00 | -0.13 |
| $\hat{\beta}_2$ | 84.22 | 12.29 | 0.001 | 0.9996 | 465.61 | 8.67 | 0.00 | 0.00 | -0.22 | -0.14 |
| $\hat{\alpha}_1 + \hat{\beta}_2$ | 86.10 | 14.17 | 0.0004 | 1.00 | 1193.03 | 8.63 | 0.00 | -0.68 | -0.17 | -0.31 |

**Table 5:** all models for predicting the relative decrease in ESS from the historical precipitation to the ESS at 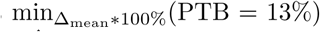; each uses a linear combination of at least one of the following: 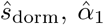 and 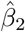 See Table 3 for variable interpretations.

| Model | AICc | $\Delta_i$ | $w_i$ | acc $w_i$ | ER | $L$ | $\hat{s}_{\text{dorm}}$ | $\hat{\alpha}_1$ | $\hat{\beta}_2$ | Adj-$R^2$ |
| --- | --- | --- | --- | --- | --- | --- | --- | --- | --- | --- |
| $\hat{s}_{\text{dorm}}$ | 42.48 | 0.00 | 0.49 | 0.49 | 1.00 | 1.37 | 0.61 | 0.00 | 0.00 | 0.44 |
| $\hat{s}_{\text{dorm}} + \hat{\beta}_2$ | 44.24 | 1.76 | 0.20 | 0.69 | 2.41 | 1.38 | 0.61 | 0.00 | -0.09 | 0.36 |
| $\hat{s}_{\text{dorm}} + \hat{\alpha}_1$ | 44.44 | 1.96 | 0.18 | 0.87 | 2.67 | 1.37 | 0.61 | 0.04 | 0.00 | 0.35 |
| $\hat{s}_{\text{dorm}} + \hat{\alpha}_1 + \hat{\beta}_2$ | 46.19 | 3.70 | 0.08 | 0.95 | 6.37 | 1.38 | 0.60 | 0.05 | -0.10 | 0.24 |
| $\hat{\beta}_2$ | 48.76 | 6.27 | 0.02 | 0.97 | 23.03 | 1.43 | 0.00 | 0.00 | -0.10 | -0.12 |
| $\hat{\alpha}_1$ | 48.86 | 6.38 | 0.02 | 0.99 | 24.28 | 1.43 | 0.00 | 0.06 | 0.00 | -0.14 |
| $\hat{\alpha}_1 + \hat{\beta}_2$ | 50.70 | 8.22 | 0.01 | 1.00 | 60.94 | 1.43 | 0.00 | 0.06 | -0.11 | -0.30 |

**Table 6:** all models for predicting the 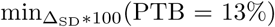; each uses a linear combination of at least one of the following predictors: 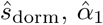, and 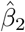. See Table 3 for variable interpretations.

| Model | AICc | $\Delta_i$ | $w_i$ | acc $w_i$ | ER | $L$ | $\hat{s}_{\text{dorm}}$ | $\hat{\alpha}_1$ | $\hat{\beta}_2$ | Adj-$R^2$ |
| --- | --- | --- | --- | --- | --- | --- | --- | --- | --- | --- |
| $\hat{s}_{\text{dorm}}$ | 56.65 | 0.00 | 0.46 | 0.46 | 1.00 | 2.79 | 1.49 | 0.00 | 0.00 | 0.50 |
| $\hat{s}_{\text{dorm}} + \hat{\beta}_2$ | 58.07 | 1.43 | 0.22 | 0.68 | 2.04 | 2.80 | 1.49 | 0.00 | -0.315 | 0.458 |
| $\hat{s}_{\text{dorm}} + \hat{\alpha}_1$ | 58.37 | 1.72 | 0.19 | 0.87 | 2.37 | 2.80 | 1.49 | 0.22 | 0.00 | 0.44 |
| $\hat{s}_{\text{dorm}} + \hat{\alpha}_1 + \hat{\beta}_2$ | 59.70 | 3.06 | 0.10 | 0.97 | 4.61 | 2.81 | 1.48 | 0.25 | -0.33 | 0.38 |
| $\hat{\beta}_2$ | 63.88 | 7.23 | 0.01 | 0.98 | 37.20 | 2.93 | 0.00 | 0.00 | -0.34 | -0.11 |
| $\hat{\alpha}_1$ | 64.00 | 7.35 | 0.01 | 0.99 | 39.46 | 2.93 | 0.00 | 0.26 | 0.00 | -0.12 |
| $\hat{\alpha}_1 + \hat{\beta}_2$ | 65.66 | 9.02 | 0.01 | 1.00 | 90.84 | 2.95 | 0.00 | 0.29 | -0.36 | -0.26 |

## References

G. Ceballos and P.R. Ehrlich. Mammal population losses and the extinction crisis. Science, 296(5569):904–907, 2002.

A.D. Barnosky, N. Matzke, S. Tomiya, G.O.U. Wogan, B. Swartz, T.B. Quental, C. Marshall, J.L. McGuire, E.L. Lindsey, K.C. Maguire, et al. Has the earth’s sixth mass extinction already arrived? Nature, 471(7336):51–57, 2011.

M.C. Urban. Accelerating extinction risk from climate change. Science, 348(6234):571–573, 2015.

Intergovernmental Panel On Climate Change. Climate change 2007: The physical science basis. Agenda, 6(07):333, 2007.

W.B. Foden, S.H.M. Butchart, S.N. Stuart, J. Vié, H.R. Akçakaya, A. Angulo, L.M. DeVantier, A. Gutsche, E. Turak, L. Cao, et al. Identifying the world’s most climate change vulnerable species: a systematic trait-based assessment of all birds, amphibians and corals. PloS one, 8(6):e65427, 2013.

L. Miles, A. Grainger, and O. Phillips. The impact of global climate change on tropical forest biodiversity in Amazonia. Global Ecology and Biogeography, 13(6):553–565, 2004.

L.J. Beaumont and L. Hughes. Potential changes in the distributions of latitudinally restricted australian butterfly species in response to climate change. Global Change Biology, 8(10):954–971, 2002.

S.J. Franks, S. Sim, and A.E. Weis. Rapid evolution of flowering time by an annual plant in response to a climate fluctuation. Proceedings of the National Academy of Sciences, 104 (4):1278–1282, 2007.

H. Gonzalo-Turpin and L. Hazard. Local adaptation occurs along altitudinal gradient despite the existence of gene flow in the alpine plant species festuca eskia. Journal of Ecology, 97 (4):742–751, 2009.

S.J. Franks, J.J. Weber, and S.N. Aitken. Evolutionary and plastic responses to climate change in terrestrial plant populations. Evolutionary applications, 7(1):123–139, 2014.

M.L. Avolio and M.D. Smith. Mechanisms of selection: phenotypic differences among genotypes explain patterns of selection in a dominant species. Ecology, 94(4):953–965, 2013.

M.L. Avolio, J.M. Beaulieu, and M.D. Smith. Genetic diversity of a dominant C<sub>4</sub> grass is altered with increased precipitation variability. Oecologia, 171(2):571–581, 2013.

A. Bush, K. Mokany, R. Catullo, A. Hoffmann, V. Kellermann, C. Sgro, S. McEvey, and S. Ferrier. Incorporating evolutionary adaptation in species distribution modelling reduces projected vulnerability to climate change. Ecology letters, 19(12):1468–1478, 2016.

M.V. Matz, E.A. Treml, G.V. Aglyamova, and L.K. Bay. Potential and limits for rapid genetic adaptation to warming in a great barrier reef coral. PLoS genetics, 14(4):e1007220, 2018.

P.H.J. Stabeno, N.A. Bond, N.B. Kachel, S.A. Salo, and J.D. Schumacher. On the temporal variability of the physical environment over the south-eastern bering sea. Fisheries Oceanography, 10(1):81–98, 2001.

N.J. Mantua, S.R. Hare, Y. Zhang, J.M. Wallace, and R.C. Francis. A pacific interdecadal climate oscillation with impacts on salmon production. Bulletin of the american Meteorological Society, 78(6):1069–1080, 1997.

J.R. Bernhardt, M.I. O’Connor, J.M. Sunday, and A. Gonzalez. Life in fluctuating environments. Philosophical Transactions of the Royal Society B, 375(1814):20190454, 2020.

F. Menu and D. Debouzie. Coin-flipping plasticity and prolonged diapause in insects: example of the chestnut weevil curculio elephas (coleoptera: Curculionidae). Oecologia, 93 (3):367–373, 1993.

B.N. Danforth. Emergence dynamics and bet hedging in a desert bee, Perdita portalis. Proceedings of the Royal Society of London. Series B: Biological Sciences, 266(1432):1985–1994, 1999.

D. Cohen. Optimizing reproduction in a randomly varying environment. Journal of Theoretical Biology, 12(1):119–129, 1966.

S.P. Ellner, N.G. Hairston Jr, C.M. Kearns, and D. Babaï. The roles of fluctuating selection and long-term diapause in microevolution of diapause timing in a freshwater copepod. Evolution, 53(1):111–122, 1999.

J. Seger. What is bet-hedging? Oxford Surveys in Evolutionary Biology, 4:182–211, 1987.

S. Ellner. Competition and dormancy: a reanalysis and review. The American Naturalist, 130(5):798–803, 1987.

D.L. Venable. Bet hedging in a guild of desert annuals. Ecology, 88(5):1086–1090, 2007.

M. Koornneef, L. Bentsink, and H. Hilhorst. Seed dormancy and germination. Current opinion in plant biology, 5(1):33–36, 2002.

E.A. Klupczyńska and T.A. Pawlowski. Regulation of seed dormancy and germination mechanisms in a changing environment. International Journal of Molecular Sciences, 22(3):1357, 2021.

K. Tielborger, M. Petruu, and C. Lampei. Bet-hedging germination in annual plants: a sound empirical test of the theoretical foundations. Oikos, 121(11):1860–1868, 2012.

P.S. Allen and S.E. Meyer. Ecology and ecological genetics of seed dormancy in downy brome. Weed Science, 50(2):241–247, 2002.

A. Carta, G. Bedini, J.V. Müller, and R.J. Probert. Comparative seed dormancy and germination of eight annual species of ephemeral wetland vegetation in a mediterranean climate. Plant Ecology, 214(2):339–349, 2013.

J.L. Walck, J.M. Baskin, C.C. Baskin, and S.N. Hidayati. Defining transient and persistent seed banks in species with pronounced seasonal dormancy and germination patterns. Seed Science Research, 15(3):189–196, 2005.

A. Sih, M.C.O. Ferrari, and D.J. Harris. Evolution and behavioural responses to human-induced rapid environmental change. Evolutionary Applications, 4(2):367–387, 2011.

A. Sih. Understanding variation in behavioural responses to human-induced rapid environmental change: a conceptual overview. Animal Behaviour, 85(5):1077–1088, 2013.

P.H. Crowley, P.C. Trimmer, O. Spiegel, S.M. Ehlman, W.S. Cuello, and A. Sih. Predicting habitat choice after rapid environmental change. The American Naturalist, 193(5):619–632, 2019.

K. Tielbörger, M.C. Bilton, J. Metz, J. Kigel, C. Holzapfel, E. Lebrija-Trejos, I. Konsens, H. A. Parag, and M. Sternberg. Middle-eastern plant communities tolerate 9 years of drought in a multi-site climate manipulation experiment. Nature communications, 5(1): 1–9, 2014.

S.J. Franks and A.E. Weis. A change in climate causes rapid evolution of multiple life-history traits and their interactions in an annual plant. Journal of evolutionary biology, 21(5): 1321–1334, 2008.

S.E. Sultan, T. Horgan-Kobelski, L.M. Nichols, C.E. Riggs, and R.K. Waples. A resurrection study reveals rapid adaptive evolution within populations of an invasive plant. Evolutionary Applications, 6(2):266–278, 2013.

J.T. Anderson and B. Song. Plant adaptation to climate change—where are we? Journal of Systematics and Evolution, 58(5):533–545, 2020.

A.J. Miller-Rushing and R.B. Primack. Global warming and flowering times in Thoreau’s concord: a community perspective. Ecology, 89(2):332–341, 2008.

B. Rhoné, R. Vitalis, I. Goldringer, and I. Bonnin. Evolution of flowering time in experimental wheat populations: a comprehensive approach to detect genetic signatures of natural selection. Evolution: International Journal of Organic Evolution, 64(7):2110–2125, 2010.

R. I. Colautti and S.C.H. Barrett. Rapid adaptation to climate facilitates range expansion of an invasive plant. Science, 342(6156):364–366, 2013.

E. Nevo, Y. Fu, T. Pavlicek, S. Khalifa, M. Tavasi, and A. Beiles. Evolution of wild cereals during 28 years of global warming in israel. Proceedings of the National Academy of Sciences, 109(9):3412–3415, 2012.

J.T. Anderson, D.W. Inouye, A.M. McKinney, R.I. Colautti, and T. Mitchell-Olds. Phenotypic plasticity and adaptive evolution contribute to advancing flowering phenology in response to climate change. Proceedings of the Royal Society B: Biological Sciences, 279 (1743):3843–3852, 2012.

S.J. Franks, N.C. Kane, N.B. O’Hara, S. Tittes, and J.S. Rest. Rapid genome-wide evolution in brassica rapa populations following drought revealed by sequencing of ancestral and descendant gene pools. Molecular Ecology, 25(15):3622–3631, 2016.

M.J. Clauss and D.L. Venable. Seed germination in desert annuals: an empirical test of adaptive bet hedging. The American Naturalist, 155(2):168–186, 2000.

S. Volis and G. Bohrer. Joint evolution of seed traits along an aridity gradient: seed size and dormancy are not two substitutable evolutionary traits in temporally heterogeneous environment. New Phytologist, 197(2):655–667, 2013.

E. Fernández-Pascual, B. Jiménez-Alfaro, J. Caujapé-Castells, R. Jaén-Molina, and T.E. Díaz. A local dormancy cline is related to the seed maturation environment, population genetic composition and climate. Annals of botany, 112(5):937–945, 2013.

C. Potvin and D. Tousignant. Evolutionary consequences of simulated global change: genetic adaptation or adaptive phenotypic plasticity. Oecologia, 108(4):683–693, 1996.

D.A. Springate, N. Scarcelli, J. Rowntree, and P.X. Kover. Correlated response in plasticity to selection for early flowering in arabidopsis thaliana. Journal of Evolutionary Biology, 24(10):2280–2288, 2011.

B.A. Robertson, J.S. Rehage, and A. Sih. Ecological novelty and the emergence of evolutionary traps. Trends in Ecology & Evolution, 28(9):552–560, 2013.

A.M. Wilczek, M.D. Cooper, T.M. Korves, and J. Schmitt. Lagging adaptation to warming climate in arabidopsis thaliana. Proceedings of the National Academy of Sciences, 111(22): 7906–7913, 2014.

J.T. Anderson and S.M. Wadgymar. Climate change disrupts local adaptation and favours upslope migration. Ecology letters, 23(1):181–192, 2020.

S. Kimball, A.L. Angertand T.E. Huxman, and D.L. Venable. Contemporary climate change in the Sonoran Desert favors cold-adapted species. Global Change Biology, 16(5):1555–1565, 2010.

T.E. Huxman, S. Kimball, A.L. Angert, J.R. Gremer, G.A. Barron-Gafford, and D.L. Venable. Understanding past, contemporary, and future dynamics of plants, populations, and communities using sonoran desert winter annuals. American Journal of Botany, 100(7): 1369–1380, 2013.

J.R. Cox, J.H. Fourie, N.F.G. Rethman, D.G. Wilcox, et al. The influence of climate and soils on the distribution of four african grasses. 1988.

A. Búrquez, A. Martínez-Yrízar, R.S. Felger, and D. Yetman. Vegetation and habitat diversity at the southern edge of the Sonoran Desert. Ecology of Sonoran Desert plants and plant communities, pages 36–67, 1999.

J.R. Gremer, S. Kimball, A.L. Angert, D.L. Venable, and T.E. Huxman. Variation in photosynthetic response to temperature in a guild of winter annual plants. Ecology, 93(12): 2693–2704, 2012.

J.R. Gremer and D.L. Venable. Bet hedging in desert winter annual plants: optimal germination strategies in a variable environment. Ecology Letters, 17(3):380–387, 2014.

J.R. Gremer, S. Kimball, and D.L. Venable. Within-and among-year germination in Sonoran Desert winter annuals: bet hedging and predictive germination in a variable environment. Ecology Letters, 19(10):1209–1218, 2016.

W.S. Cuello, J.R. Gremer, P.C. Trimmer, A. Sih, and S.J. Schreiber. Predicting evolutionarily stable strategies from functional responses of Sonoran Desert annuals to precipitation. Proceedings of the Royal Society B, 286(1894):20182613, 2019.

R.C. Brusca, J.F. Wiens, W.M. Meyer, J. Eble, K. Franklin, J.T. Overpeck, and W. Moore. Dramatic response to climate change in the southwest: Robert Whittaker’s 1963 Arizona Mountain plant transect revisited. Ecology and Evolution, 3(10):3307–3319, 2013.

A.G. Pendergrass, R. Knutti, F. Lehner, C. Deser, and B.M. Sanderson. Precipitation variability increases in a warmer climate. Scientific Reports, 7(1):17966, 2017.

F. Dominguez, E. Rivera, D.P. Lettenmaier, and C.L. Castro. Changes in winter precipitation extremes for the western United States under a warmer climate as simulated by regional climate models. Geophysical Research Letters, 39(5), 2012.

P.Y. Groisman and R.W. Knight. Prolonged dry episodes over the conterminous united states: New tendencies emerging during the last 40 years. Journal of Climate, 21(9): 1850–1862, 2008.

R. James and R. Washington. Changes in African temperature and precipitation associated with degrees of global warming. Climatic change, 117(4):859–872, 2013.

K.E. Trenberth, A. Dai, G. Van Der Schrier, P.D. Jones, J. Barichivich, K.R. Briffa, and J. Sheffield. Global warming and changes in drought. Nature Climate Change, 4(1):17–22, 2014.

G.M. MacDonald. Water, climate change, and sustainability in the southwest. Proceedings of the National Academy of Sciences, 107(50):21256–21262, 2010.

Amy L Angert, Jonathan L Horst, Travis E Huxman, and D Lawrence Venable. Phenotypic plasticity and precipitation response in sonoran desert winter annuals. American Journal of Botany, 97(3):405–411, 2010.

T.R. Huggins, B.Al. Prigge, M.R. Sharifi, and P.W. Rundel. The effects of long-term drought on host plant canopy condition and survival of the endangered Astragalus jaegerianus (Fabaceae). Madroño, 57(2):120–128, 2010.

J.B. Bradford, D.R. Schlaepfer, W.K. Lauenroth, and K.A. Palmquist. Robust ecological drought projections for drylands in the 21st century. Global change biology, 26(7):3906–3919, 2020.

A. Dai. Drought under global warming: a review. Wiley Interdisciplinary Reviews: Climate Change, 2(1):45–65, 2011.

D.L. Venable. Long-term population dynamics of individually mapped Sonoran Desert winter annuals from the Desert Laboratory, Tucson, AZ.

C. E. Pake and D.L. Venable. Seed banks in desert annuals: implications for persistence and coexistence in variable environments. Ecology, 77(5):1427–1435, 1996.

M. Benaïm and S.J. Schreiber. Persistence of structured populations in random environments. Theoretical Population Biology, 76(1):19–34, 2009.

S.J. Schreiber and M. Benaïm. Persistence and extinction for stochastic ecological models with internal and external variables. Journal of Mathematical Biology, 79(1):393–431, 2019.

J. Mullahy. Specification and testing of some modified count data models. Journal of Econometrics, 33(3):341–365, 1986.

P.L. Chesson and S. Ellner. Invasibility and stochastic boundedness in monotonic competition models. Journal of Mathematical Biology, 27(2):117–138, 1989.

Y. Bai, T.A. Scott, and Q. Min. Climate change implications of soil temperature in the Mojave Desert, USA. Frontiers of Earth Science, 8(2):302–308, 2014.

R.E. Symonds and A. Moussalli. A brief guide to model selection, multimodel inference and model averaging in behavioural ecology using akaike’s information criterion. Behavioral Ecology and Sociobiology, 65(1):13–21, 2011.

K.P. Burnham and D.R. Anderson. Sociological methods & research. Sociological Methods Research, 33(2):261–304, 2004.

K.P. Burnham, D.R. Anderson, and K.P. Huyvaert. Aic model selection and multimodel inference in behavioral ecology: some background, observations, and comparisons. Behavioral Ecology and Sociobiology, 65(1):23–35, 2011.

S.C. Choi. Tests of equality of dependent correlation coefficients. Biometrika, 64(3):645–647, 1977.

E.C. Fieller, H.O. Hartley, and E.S. Pearson. Tests for rank correlation coefficients. i. Biometrika, 44(3/4):470–481, 1957.

D.R. Easterling, G.A. Meehl, C. Parmesan, S.A. Changnon, T.R. Karl, and L.O. Mearns. Climate extremes: observations, modeling, and impacts. science, 289(5487):2068–2074, 2000.

J.F. McLaughlin, J.J. Hellmann, C.L. Boggs, and P.R. Ehrlich. Climate change hastens population extinctions. Proceedings of the National Academy of Sciences, 99(9):6070–6074, 2002.

S. Vincenzi. Extinction risk and eco-evolutionary dynamics in a variable environment with increasing frequency of extreme events. Journal of The Royal Society Interface, 11(97): 20140441, 2014.

D.L. Venable, C.E. Pake, and A.C. Caprio. Diversity and coexistence of sonoran desert winter annuals. Plant Species Biology, 8(2-3):207–216, 1993.

F. Shreve and I.L. Wiggins. Vegetation and flora of the Sonoran Desert, volume 591. Stanford University Press, 1964.

T. Philippi. Bet-hedging germination of desert annuals: beyond the first year. The American Naturalist, 142(3):474–487, 1993.

S. Ellner. ESS germination strategies in randomly varying environments. I. Logistic-type models. Theoretical Population Biology, 28(1):50–79, 1985.

D. Cohen. A general model of optimal reproduction in a randomly varying environment. The Journal of Ecology, pages 219–228, 1968.

A.L. Angert, T.E. Huxman, G.A. Barron-Gafford, K.L. Gerst, and D.L. Venable. Linking growth strategies to long-term population dynamics in a guild of desert annuals. Journal of Ecology, 95(2):321–331, 2007.

T.E. Huxman, G.A. Barron-Gafford, K.L. Gerst, A.L. Angert, A.P. Tyler, and D.L. Venable. Photosynthetic resource-use efficiency and demographic variability in desert winter annual plants. Ecology, 89(6):1554–1563, 2008.

M.K.J. Ooi. Seed bank persistence and climate change. Seed Science Research, 22(S1): S53–S60, 2012.

F.S. Chapin III, K. Autumn, and F. Pugnaire. Evolution of suites of traits in response to environmental stress. The American Naturalist, 142:S78–S92, 1993.

A.L. Angert, S. Kimball, M. Peterson, T.E. Huxman, and D.L. Venable. Phenotypic constraints and community structure: Linking trade-offs within and among species. Evolution, 68(11):3149–3165, 2014.

S. Kimball, J.R. Gremer, A.L. Angert, T.E. Huxman, and D.L. Venable. Fitness and physiology in a variable environment. Oecologia, 169(2):319–329, 2012.

I. Noy-Meir. Desert ecosystems: environment and producers. Annual review of ecology and systematics, 4(1):25–51, 1973.

T.R. Ault, J.S. Mankin, B.I. Cook, and J.E. Smerdon. Relative impacts of mitigation, temperature, and precipitation on 21st-century megadrought risk in the american southwest. Science Advances, 2(10):e1600873, 2016.

Ming Liu, Dustin R Rubenstein, Wei-Chung Liu, and Sheng-Feng Shen. A continuum of biological adaptations to environmental fluctuation. Proceedings of the Royal Society B, 286(1912):20191623, 2019.

D.L. Venable and J.S. Brown. The selective interactions of dispersal, dormancy, and seed size as adaptations for reducing risk in variable environments. The American Naturalist, 131(3):360–384, 1988.

J.R. Gremer, S. Kimball, K.R. Keck, T.E. Huxman, A.L. Angert, and D.L. Venable. Water-use efficiency and relative growth rate mediate competitive interactions in sonoran desert winter annual plants. American Journal of Botany, 100(10):2009–2015, 2013.

R.T. Corlett and D.A. Westcott. Will plant movements keep up with climate change? Trends in Ecology & Evolution, 28(8):482–488, 2013.

A.S. Jump and J. Penuelas. Running to stand still: adaptation and the response of plants to rapid climate change. Ecology letters, 8(9):1010–1020, 2005.

R. Bertrand, J. Lenoir, C. Piedallu, G. Riofrio-Dillon, P. de Ruffray, C. Vidal, J. Pierrat, and J. Gégout. Changes in plant community composition lag behind climate warming in lowland forests. Nature, 479(7374):517–520, 2011.

E. Hamann, A.E. Weis, and S.J. Franks. Two decades of evolutionary changes in brassica rapa in response to fluctuations in precipitation and severe drought. Evolution, 72(12): 2682–2696, 2018.

E.E. Dickman, L.K. Pennington, S.J. Franks, and J.P. Sexton. Evidence for adaptive responses to historic drought across a native plant species range. Evolutionary Applications, 12(8):1569–1582, 2019.

J. Metz, C. Lampei, L. Bäumler, H. Bocherens, H. Dittberner, L. Henneberg, J. de Meaux, and K. Tielbörger. Rapid adaptive evolution to drought in a subset of plant traits in a large-scale climate change experiment. Ecology Letters, 23(11):1643–1653, 2020.

A.P. Hendry. Eco-evolutionary dynamics. Princeton university press, 2016.

R.G. Shaw and J.R. Etterson. Rapid climate change and the rate of adaptation: insight from experimental quantitative genetics. New Phytologist, 195(4):752–765, 2012.

A. Loydi and S.L. Collins. Extreme drought has limited effects on soil seed bank composition in desert grasslands. Journal of Vegetation Science, 32(5):e13089, 2021.

M.L. LaForgia, M.J. Spasojevic, E.J. Case, A.M. Latimer, and S.P. Harrison. Seed banks of native forbs, but not exotic grasses, increase during extreme drought. Ecology, 99(4): 896–903, 2018.

Alan R Templeton and Donald A Levin. Evolutionary consequences of seed pools. The American Naturalist, 114(2):232–249, 1979.

Q. Guo, P.W. Rundel, and D.W. Goodall. Horizontal and vertical distribution of desert seed banks: patterns, causes, and implications. Journal of arid environments, 38(3):465–478, 1998.

J.L. Horst and D.L. Venable. Frequency-dependent seed predation by rodents on Sonoran Desert winter annual plants. Ecology, 2017.

L.F. Soholt. Consumption of primary production by a population of kangaroo rats (dipodomys merriami) in the mojave desert. Ecological Monographs, 43(3):357–376, 1973.

J.S. Brown, B.P. Kotler, R.J. Smith, and W.O. Wirtz. The effects of owl predation on the foraging behavior of heteromyid rodents. Oecologia, 76(3):408–415, 1988.

K.J. Iknayan and S.R. Beissinger. Collapse of a desert bird community over the past century driven by climate change. Proceedings of the National Academy of Sciences, 115(34): 8597–8602, 2018.

E. A. Riddell, K.J. Iknayan, B.O. Wolf, B. Sinervo, and S.R. Beissinger. Cooling requirements fueled the collapse of a desert bird community from climate change. Proceedings of the National Academy of Sciences, 116(43):21609–21615, 2019.

S.R. Archer and K.I. Predick. Climate change and ecosystems of the southwestern united states. Rangelands, 30(3):23–28, 2008.

M.K.J. Ooi, T.D. Auld, and A.J. Denham. Climate change and bet-hedging: interactions between increased soil temperatures and seed bank persistence. Global Change Biology, 15(10):2375–2386, 2009.

J. Kochanek, Y.M. Buckley, R.J. Probert, S.W. Adkins, and K.J. Steadman. Pre-zygotic parental environment modulates seed longevity. Austral Ecology, 35(7):837–848, 2010.

S.E. Newson, S. Mendes, H.Q.P. Crick, N.K. Dulvy, J.D.R. Houghton, G.C. Hays, A.M. Hutson, C.D. MacLeod, G.J. Pierce, and R.A. Robinson. Indicators of the impact of climate change on migratory species. Endangered Species Research, 7(2):101–113, 2009.

P. Pliscoff, F. Luebert, H.H. Hilger, and A. Guisan. Effects of alternative sets of climatic predictors on species distribution models and associated estimates of extinction risk: A test with plants in an arid environment. Ecological Modelling, 288:166–177, 2014.

M.C. Jones, S.R. Dye, J.A. Fernandes, T.L. Frölicher, J.K. Pinnegar, R. Warren, and W. Cheung. Predicting the impact of climate change on threatened species in UK waters. PloS one, 8(1):e54216, 2013.

H.M. Pereira, P.W. Leadley, V. Proença, R. Alkemade, J.P.W. Scharlemann, J.F. Fernandez-Manjarrés, M.B. Araújo, P. Balvanera, R. Biggs, W.W.L. Cheung, et al. Scenarios for global biodiversity in the 21st century. Science, 330(6010):1496–1501, 2010.

A.A. Hoffmann and C.M. Sgrò. Climate change and evolutionary adaptation. Nature, 470 (7335):479–485, 2011.

K.P. Burnham and D.R. Anderson. Model Selection and Multimodel Inference: A practical Information-theoretic Approach. 2002.

P. M. Lukacs, K.P. Burnham, and D.R. Anderson. Model selection bias and freedman’s paradox. Annals of the Institute of Statistical Mathematics, 62(1):117, 2010.

